# Hierarchical Network Model Excitatory-Inhibitory Tone Shapes Alternative Strategies for Different Degrees of Uncertainty in Multi-Attribute Decisions

**DOI:** 10.1101/2020.01.28.923649

**Authors:** Warren Woodrich Pettine, Kenway Louie, John D Murray, Xiao Jing Wang

## Abstract

We investigated two-attribute, two-alternative decision-making in a hierarchical neural network with three layers: an input layer encoding choice alternative attribute values; an intermediate layer of modules processing separate attributes; and a choice layer producing the decision. Depending on intermediate layer excitatory-inhibitory (E/I) tone, the network displays three distinct regimes characterized by linear (I), convex (II) or concave (III) choice indifference curves. In regimes I and II, each option’s attribute information is additively integrated. To maximize reward at low environmental uncertainty, the system should operate in regime I. At high environmental uncertainty, reward maximization is achieved in regime III, with each attribute module selecting a favored alternative, and the ultimate decision based upon comparison between outputs of attribute processing modules. We then use these principles to examine multi-attribute decisions with autism-related deficits in E/I balance, leading to predictions of different choice patterns and overall performance between autism and neurotypicals.

## 1 Introduction

We are constantly faced with decisions between alternatives defined by multiple attributes. The true value of each attribute is at times clear, and other times uncertain. For example, on Friday one might choose between main courses at a restaurant where the flavor or healthiness attributes of all the dishes are familiar. The following Wednesday might be at a restaurant with an unknown cuisine, where one is highly uncertain as to different items’ flavor or healthiness. To ensure the best meal, the brain must be able to optimize choice in both environments.

Systems neuroscientists have, for many years, been studying the specific circuits engaged in this kind of multi-attribute decision-making. Based on a robust set of electrophysiology and imaging findings (Xie and Padoa-Schioppa 2016; Raghuraman and Padoa-Schioppa 2014; Padoa-Schioppa and Assad 2006; O’Neill and Schultz 2018; Morrison and Salzman 2009; Conen and Padoa-Schioppa 2015; Chib et al. 2009; Pastor-Bernier, Stasiak, and Schultz 2019), many hold that all attribute signals are available in brain areas proximal to the final decision (Levy and Glimcher 2012; Padoa-Schioppa and Conen 2017). Indeed, when attribute values are clear, multi-attribute choice theoretically is simple: linearly weight and combine all attributes associated with a choice alternative, then select the one with the larger value. Though the subjective value of an attribute might be non-linearly related to the quantity offered, when the final choice is made in an environment without uncertainty, a weighted linear combination of attributes optimizes the choice between options (Nicholson and Snyder 2007).

However, an alternative perspective has recently arisen that takes into account the multiple brain areas implicated in decisions (Wunderlich, Rangel, and O’Doherty 2009; Peck, Lau, and Salzman 2013; Paton et al. 2006; Munuera, Rigotti, and Salzman 2018; Louie and Glimcher 2010; Chen and Stuphorn 2015; Steinmetz et al. 2019). Under this alternate view, a signal is transformed as it moves from hierarchically lower areas to those that are proximal to the final decision (Cisek and Kalaska 2010; Cisek 2012; Yoo and Hayden 2018). Such a process can allow for parallel computations, and produce clearer separation between the representation of choice alternatives where the decision is reached. Furthermore, these transformations can be highly non-linear, providing an additional layer of flexibility in the decision process.

When investigating the non-linear transformations that can occur in populations of neurons, it is imperative to consider excitatory and inhibitory (E/I) tone. In sensory areas, E/I tone can shape the stimulus tuning curve and responses timing (Mariño et al. 2005; Wehr and Zador 2003). When sensory information is used in perceptual decision-making, E/I tone can dictate the trade-off between speed and accuracy (Wong and Wang, 2006). In working memory tasks, E/I tone defines a network’s ability to maintain a memory, and that memory’s susceptibility to a distractor (Brunel and Wang 2001; Compte et al. 2000). When multiple areas interact, the E/I tone can define their function in either memory maintenance or decision-making (Murray, Jaramillo, and Wang 2017). On a more theoretical level, E/I tone can shape a system’s response as an input signal increases (Ahmadian and Miller 2019) and its transition to chaotic dynamics (Vreeswijk and Sompolinsky 1998). Yet, it remains to be described how E/I tone defines the functional interaction and transformation of multiple signals passing through a brain area, as in a hierarchical system.

Though a prior biophysical model of hierarchical neural networks engaged in multi-attribute decisionmaking captured key features of human reaction times and functional imaging (Hunt, Dolan, and Behrens 2014), how a hierarchical system shapes multi-attribute decisions is largely a matter of conjecture (Hunt and Hayden 2017). By their very nature, these transformations distort the signals available for the final decision. Distributed models have been shown to benefit motor control (Christopoulos, Bonaiuto, and Andersen 2015; Cisek 2006), but in what situations might such transformations improve a decision process?

To approach this question, we examine how E/I tone in hierarchical networks governs multi-attribute decision-making under varying degrees of environmental uncertainty. It is often the case that noise in the environment creates uncertainty as to the true value of attributes (Bach and Dolan 2012). Additionally, environmental uncertainty offers an entry-point to understanding neuropsychiatric conditions, such as autism spectrum disorder (ASD). A myriad of findings suggest that the diagnostic signs of behavioral rigidity and sensory abnormalities (American Psychiatric Association 2013) may be related to an intolerance of environmental uncertainty (Foss-Feig et al. 2017; Fujino et al. 2017; Boulter et al. 2014; Van de Cruys et al. 2014; Vasa et al. 2018). A mechanistic understanding of how hierarchical neural systems handle environmental uncertainty when making decisions can not only provide knowledge as to how the brain functions, but also may produce hypotheses as to the etiology of ASD.

We find that the E/I tone of the network creates distinct regimes defined by their choice indifference curve: a linear weighting of attribute values (regime I); a convex preference for balanced attributes (regime II); or a concave increased weighting of the larger attribute value (regime III). We then show that the degree of reward maximization by these regimes depends on the level of environmental uncertainty. When uncertainty is low, regimes I or II - where the choice area has access to all signal information - achieve the greatest reward. However, when uncertainty is high, regime III - with strong transforms along the hierarchy favoring the larger attribute value - on average, maximizes reward. After a detailed investigation of how these regimes arise and why they are optimal, we hypothesize a framework where the E/I tone can be modified according to the level of uncertainty. The framework is then used to create a model of neurotypicals with the full range of E/I tone, and a model of ASD populations defined by limited inhibitory tone. Based on the models, we predict that neurotypical and ASD subjects will adopt a linear weighting of attributes in environments with low uncertainty (regimes I/II). As environmental uncertainty increases, neurotypical subjects will fully move into regime III, while ASD individuals will not. This will result in larger performances differences at high environmental uncertainty levels.

Using a hierarchical network parameterized by E/I tone, we are able to show that the the network can adopt multiple regimes (including linear) to maximize reward in various environments, and then we use these findings to provide an etiological hypothesis for behavior observed in ASD.

## 2 Results

### 2.1 Networks Perform Multi-Attribute Decision Task Under Uncertainty

To investigate the behavior of neural networks in environments with varying degrees of uncertainty, we used a multi-attribute decision task with varying degrees of environmental noise. On any given trial, the neural networks were presented with two choice alternatives (*A* or *B*). These choice alternatives were each composed of two attributes (1 or 2). For simplicity, attributes were assumed to have equal contributions to subjective value and actions were assumed to be symmetric. When indexing, the first value refers to the choice alternative, and the second to the attribute. Thus, choice alternative *A* was presented to the network using input *I*_*A*,1_ and input *I*_*A*,2_, while choice alternative *B* was presented to the network using input *I*_*B*,1_ and input *I*_*B*,2_. Inputs were translated from offer values to firing rates. As the final choice area competition was between alternatives, the signal at that stage is denoted by *C_A_* or *C_B_*, where the subscript indicates the choice alternative. On a given trial, the choice between *A* or *B* was determined by the first population in the choice area (*C_A_* or *C_B_*) to cross a firing rate threshold (35 Hz). To control the level of uncertainty in the environment, we added a noise term *η_I_* to each input at the beginning of the trial, independently drawn from a normal distribution centered on 0, with a standard deviation of *σ_η_I__*.

We created two basic network frameworks: the Linear Network and the Hierarchical Network. Areas in these networks were composed of mean-field approximations of population firing rates (see Methods). As the name suggests, the Linear Network computed an exact linear weighted sum of the attributes (weight = 0.5). Thus, with the Linear Network, the choice area received a signal that was linearly translated from the presented attributes of the choice alternatives (Figure 1A). This network has a similar architecture to that presented in (Rustichini and Padoa-Schioppa 2015), and represents the case where all attribute signals are fully available to a choice area.

**Figure 1:**
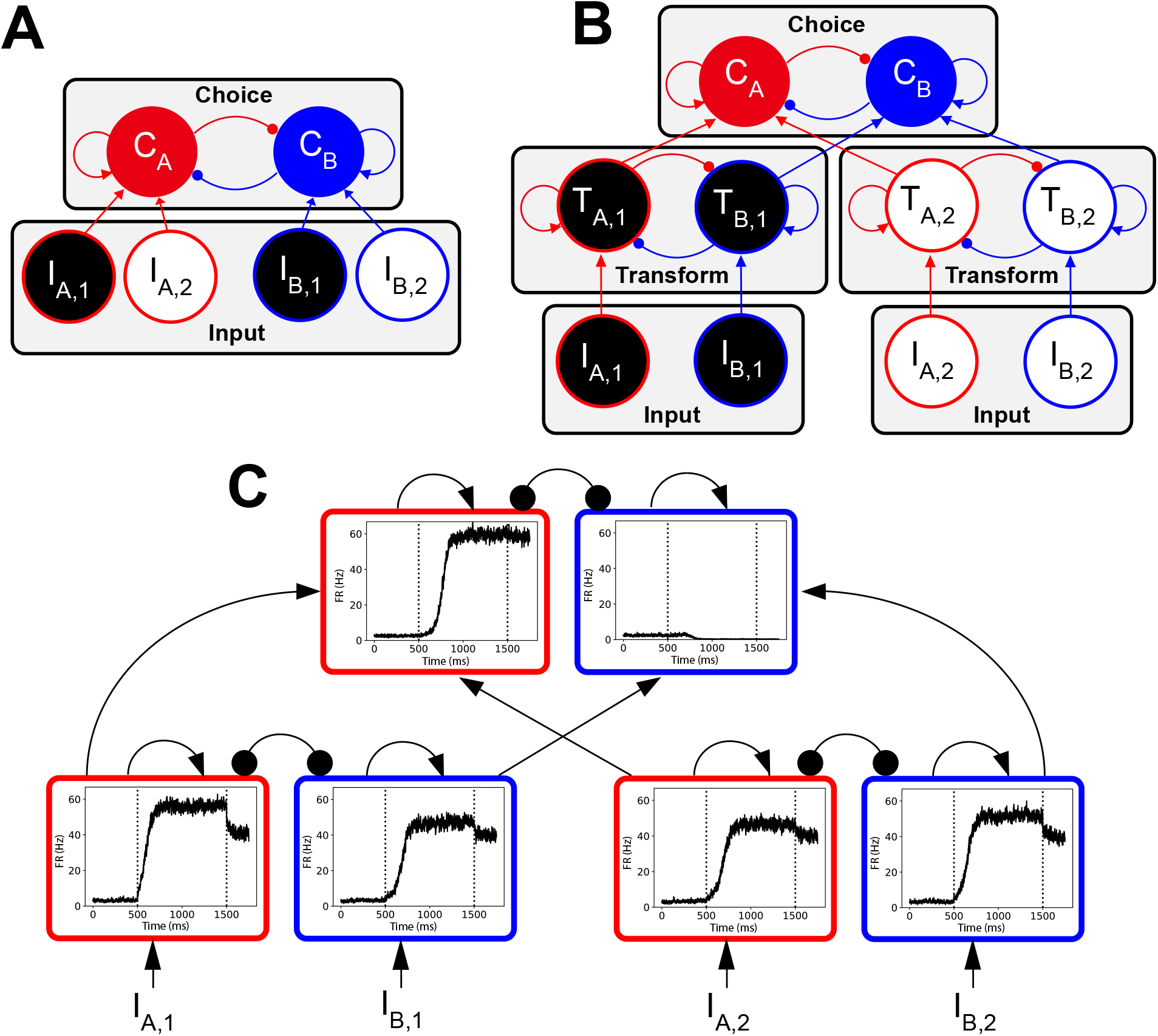
Network Schematics and Sample Trial. *I*: input; *C* choice layer synaptic gating variable; *T*: intermediate transform layer synaptic gating variable; red: choice *A*; blue: choice *B*; black: attribute 1; white: 2; arrows: excitation; circles: inhibition. Both networks include an input layer and a final choice layer. (A) The Linear Network consists of two layers, with attribute signals from the input layer directly transmitted to the final choice layer. (B) The Hierarchical Network includes an additional intermediate layer that performs a functional transformation on the attribute signals prior to their passing to the choice area. (C) A sample trial of the Hierarchical Network, with intermediate layer weights *J*_+_ = 0.34 nA and *J*_−_ = −0.02. For each area, the X-axis indicates time, while the Y-axis indicates the population firing rate. The first vertical line indicates the onset of the offer value signal, and the second vertical line indicates termination of the offer value signal.

In the Hierarchical Network (Figure 1B), inputs were first transformed by intermediate, attribute-specific areas. The intermediate layer transforming the input signal was a nonlinear dynamical system, composed of attractor states associated with each choice alternative (see Methods). To denote the transformation performed by an intermediate layer area on the input signal, its output signal will be referred to as *T*, such that *I*_*A*,1_ is related to *T*_*A*,1_, etc. These continuous output signals were then fed into a choice area that determined the decision on a given trial. A single trial of the Hierarchical Network is shown in Figure 1C. This framework, similar in structure to that presented in (Hunt, Dolan, and Behrens 2014), represents the simplest form of a network with parallel processing streams, where transformations are performed on attribute signals passing through specialized areas prior to the final decision.

To first verify that the networks were able to perform the task, we computed simple psychometric functions of model choice behavior. This was done by varying the input *I*_*A*,1_, while keeping inputs *I*_A,2_, *I*_*B*,1_, and *I*_*B*,2_ fixed. We then measured the proportion of trials that networks chose *A*, and fit a sigmoid function to the resultant *P*(*A*) (see Methods). We did this for environments where there was either no uncertainty (*σ_η_I__* = 0 Hz), or a moderate amount of uncertainty (*σ_η_I__* = 0.75 Hz).

Figure 2A shows the psychometric functions for the Linear Network in a certain environment (magenta) and an uncertain environment (green). The same is shown in Figure 2B for the Hierarchical Network with intermediate layer recurrent excitation (*J*_+_) = 0.33 nA and cross inhibition (*J*_−_) = −0.03 nA. Both networks generated expected psychometric functions, with larger input value differences resulting in better performance. Furthermore, the slope of the functions was, as anticipated, determined by the level of environmental uncertainty, such that the greater uncertainty produced a less precise choice performance.

**Figure 2:**
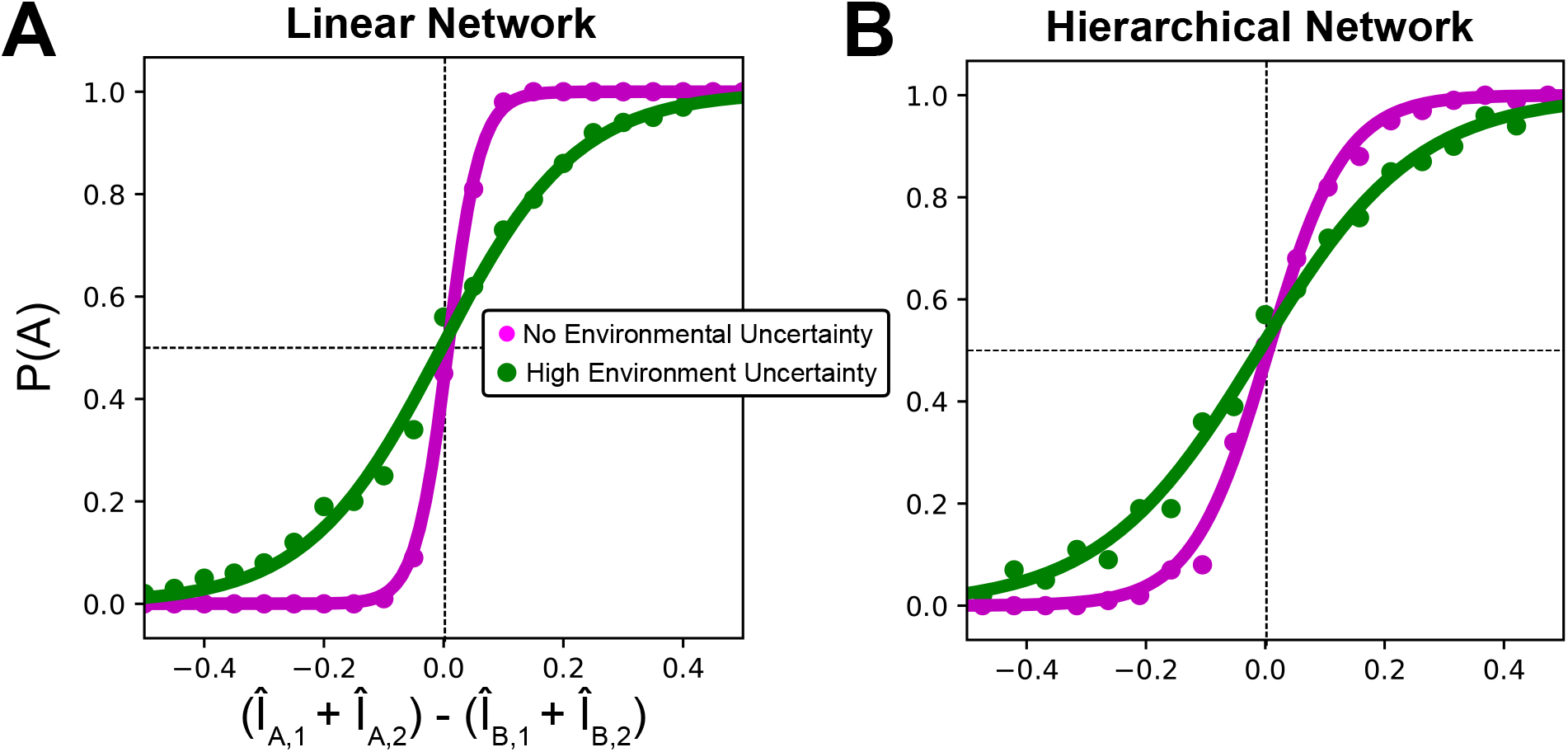
Choice Performance Under Varying Environmental Uncertainty. The proportion of trials where the network chose option *A*, *P*(*A*), as a function of the difference in offer inputs is shown, along with a fit sigmoid. Input *I*_*A*,1_ was varied as all other inputs were held fixed. On the x-axis, inputs were normalized (*Î*) by the maximum choice alternative offer value’s linear sum. (A) Performance of the Linear Network when attribute values are certain, with *σ_η_I__* = 0.0 Hz (magenta), and when attribute values are uncertain, with *σ_η_I__* = 0.75 Hz (green). (B) The same is given for the Hierarchical Network, with a intermediate layer weights of *J*_+_ = 0.33 nA and *J*_−_ = −0.03 nA.

### 2.2 The Intermediate Layer Weights of Hierarchical Networks Dictate Decision Regimes

Having verified the networks were able to perform the task, we next examined how they utilize attribute information. While the Linear Network performs a simple linear sum of the attributes, the Hierarchical Network first passes the input signals through attribute-specific areas (Figure 1B and Figure 3A). This transform of inputs is defined by the recurrent excitation and cross-inhibition of areas in the intermediate layer.

**Figure 3:**
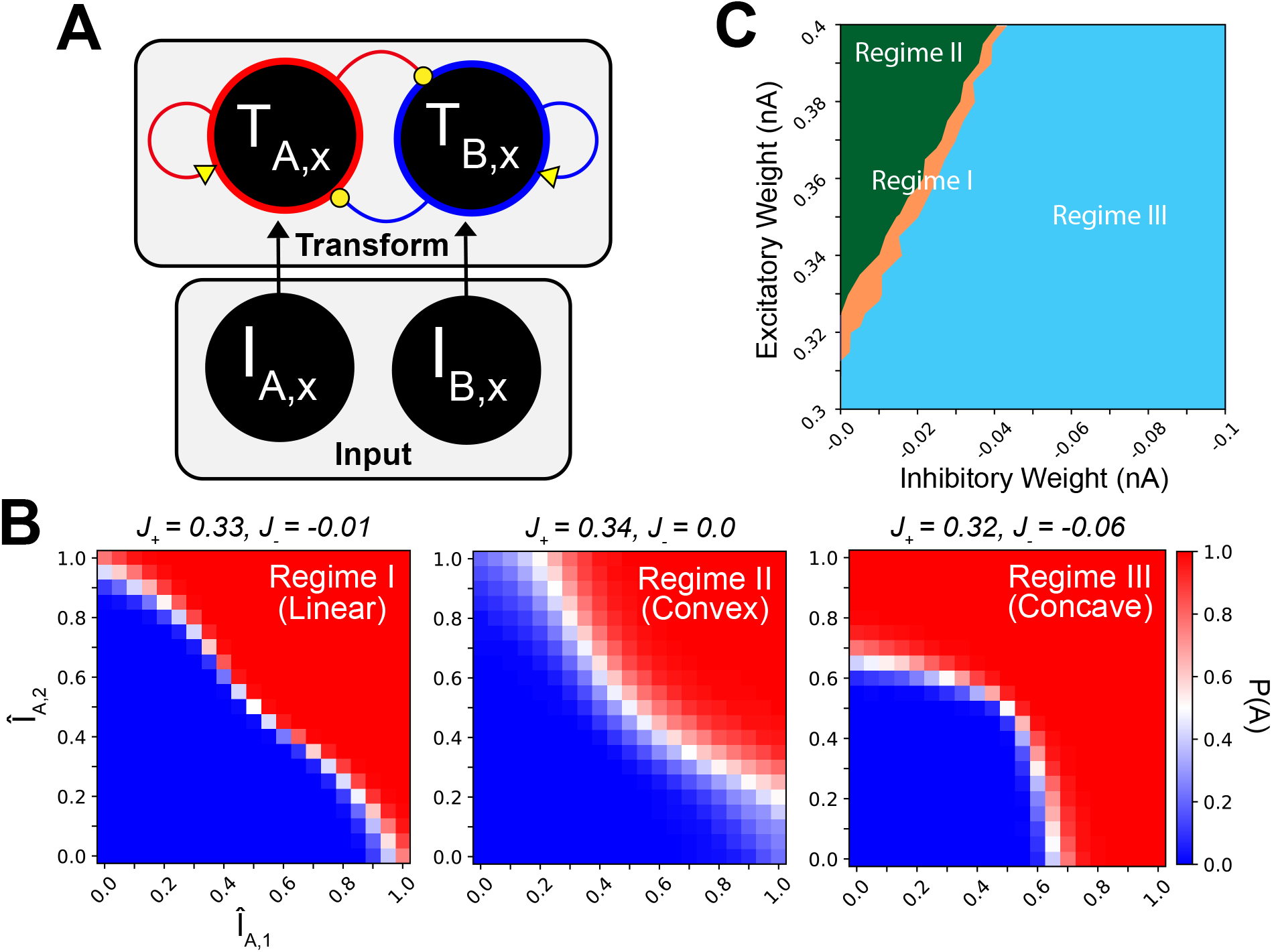
Decision Regimes of Hierarchical Networks. *I*: input; *Î*: input normalized by the reference choice alternative values’ linear sum; *T*: transform layer synaptic gating variable; *P*(*A*): proportion of 1,000 trials where alternative A was chosen; *J*_+_: excitatory weight; *J*_−_: inhibitory weight; yellow: varied weights; forest green: regime II; tan: regime I; sky blue: regime III. The first subscript indicates choice *A* or *B*, and the second subscript x indicates a generic attribute. Weights were systematically varied identically in all intermediate layer areas. (A) A diagram of a generic intermediate layer area and its inputs. (B) Examples of regime I (linear), regime II (convex) and regime III (concave) decision regimes. (C) The space of E/I weights is partitioned according to the decision regime of their indifference curves.

To investigate how the transform of attribute inputs changed with weights, we used indifference curves. When generating an indifference curve for each weight-configuration, we varied the composition and magnitude of attributes in a single choice alternative (*I*_*A*,1_ and *I*_*A*,2_), while holding the composition and magnitude of the other choice alternative (IB,1 and IB,2) constant as a reference. For some values of *I*_*A*,1_ and *I*_*A*,2_, (for example when both are much less than their counterparts) *P*(*A*) will be low. For other values of *I*_*A*,1_ and *I*_*A*,2_, (for example when both are much greater than their counterparts) *P*(*A*) will be high. However, there will be a set of values of *I*_*A*,1_ and *I*_*A*,2_ where the subject is indifferent between the choices and *P*(*A*) ≈ 0.5.

The values where *P*(*A*) ≈ 0.5 were used to fit the indifference curve.

The Linear Network by definition computes a weighted sum of the attributes, so its indifference curve is trivially linear, following the line where *I*_*A*,1_ +*I*_*A*,2_ = *I*_*B*,1_ + *I*_*B*,2_. For the Hierarchical Network, however, the indifference curve provides a tool for analyzing how the network weighs and combines attribute information (Figure 3). A linear indifference curve indicates that the network is performing a linear combination (similar to that of the Linear Network). A convex indifference curve indicates that the network places greater weight on attribute values composing a choice alternative that is balanced between the attributes, and less weight on attribute values composing an unbalanced choice alternative. A concave indifference curve indicates the opposite; specifically the network under-weighs the smaller attribute value, and instead place preferentially greater weight on the larger of the attributes.

We parametrically varied the levels of excitatory (0.30 nA to 0.40 nA, spaced 0.05 nA) and inhibitory weights (0.0 nA to −0.10 nA, spaced 0.05 nA) within intermediate layer areas. The varied weights are colored yellow in Figure 3A. For each weight-configuration, we held the inputs *I*_*B*,1_ and *I*_*B*,2_ constant at 20 Hz, while independently varying *I*_*A*,1_ and *I*_*A*,2_ from 0 to 40 Hz. We did this for 1,000 trials at each combination and from those calculated *P*(*A*).

We then fit a single-parameter exponential function to the points where *P*(*A*) ≈ 0.5 (see Methods). That single parameter determined the curvature, such that a value of 1 indicates a linear indifference curve, a value less than one indicates convex, and a value greater than 1 indicates concave. We found that at different levels of excitation and inhibition, the network adopted distinct decision regimes, corresponding to linear (regime I), convex (regime II) and concave (regime III) indifference curves (Figure 3B). The appearance of these decision regimes formed distinct regions in the weight-space, such that we were able to use these regions to partition the weight-space (Figure 3C). This revealed that by configuring the levels of excitation and inhibition in the intermediate layer, the Hierarchical Network can adopt different decision-making regimes.

### 2.3 Optimal Intermediate Layer Weights Depend on Magnitude of Environmental Uncertainty

In what settings might distinct decision regimes in a hierarchical neural system be advantageous? Indeed, if the goal is to maximize reward in an environment without uncertainty, a linear combination of attribute values achieves optimality (Nicholson and Snyder 2007). Yet, many real-world decisions are made in environments where the true value of an attribute is uncertain. For example, when examining a menu to choose a meal at a restaurant, there may be dishes containing ingredients with which one is only vaguely familiar. We therefore investigated the performance of these networks in environments where varying degrees of environmental noise created uncertainty as to the true attribute values.

To do this, we sequentially presented networks with an array of 630 choice alternative combinations, which were translated to firing rates of 10 to 20 Hz, spaced by 1 Hz. The inputs were produced combinatorially, such that on one trial the *A* attributes might be, [11 Hz, 12 Hz] and *B*, [13 Hz, 18 Hz]. On the next trial, *A* could be composed of, [11 Hz, 13 Hz] and *B*, [12 Hz, 18 Hz], etc. These choices were presented in blocks, where each block had a fixed *σ_η_I__* that ranged from 0 to 2 Hz.

For the Linear Network and each weight-configuration of the Hierarchical Network, we calculated the average amount of reward-per-trial. Heatmaps of the reward-per-trial for the Hierarchical Network are shown in Figure 4A, with the performance of the Linear Network indicated by the black dot on the heatmap colorbar. The lower-bound of the heatmap colorbar indicates the minimum possible reward-per-trial (set to 0), and the upper-bound indicates the maximum possible reward-per-trial (set to 1). Weight-configurations with regime I indifference curves are outlined in white. The region to the upper left of the white outlines corresponds to regime II, and to the right to regime III. On the left of Figure 4A is performance in a certain environment, and on the right performance in an environment with a high amount of environmental uncertainty. In both heatmaps, the optimally performing weight-configuration is indicated by a black “X.” The shift in the location of optimal weight-configuration is shown with more detail in Figure 4B, which plots the optimal excitatory weight (magenta) and inhibitory weight (turquoise) across different magnitudes of environmental uncertainty. Note that the optimal excitatory weight fluctuates in a narrow range (0.32 to 0.34 nA), while the optimal inhibitory weight magnitude increases monotonically (−0.01 to −0.1 nA). The shape of the indifference curve produced by these weights is shown in Figure 4C, where the decision regime starts in regime I (linear), and then moves to regime III, becoming increasingly concave as the environment becomes more uncertain. Thus, the Hierarchical Network displays distinct decision regimes that maximize reward under a variety of environmental uncertainty levels. Furthermore, the shift in regimes with uncertainty is primarily achieved through increasing inhibitory tone.

**Figure 4:**
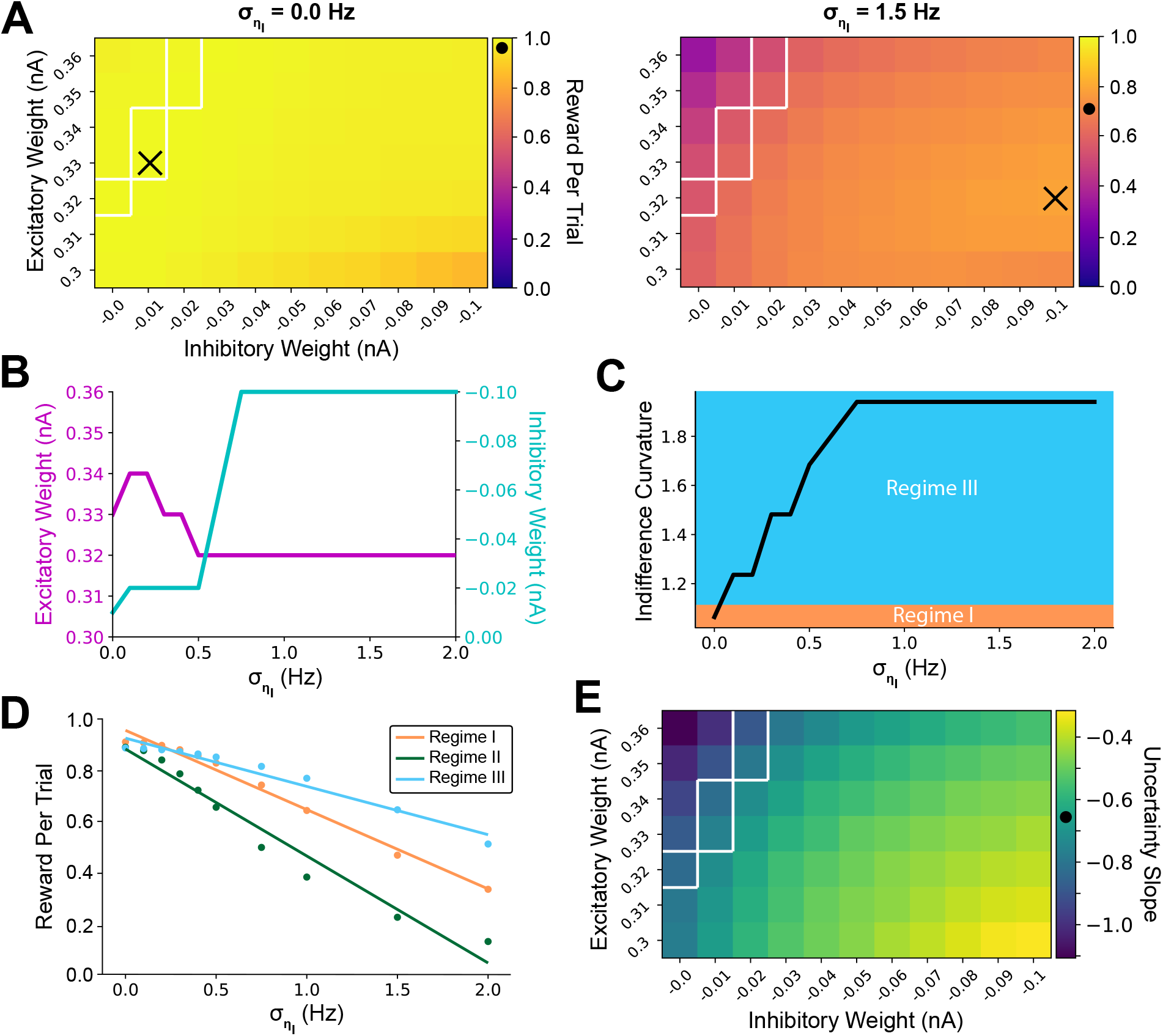
Hierarchical Network E/I Tone Governs Reward Performance Under Varying Uncertainty. (A) Heatmaps of the average reward-per-trial for different configurations of intermediate layer E/I weights. The X-axis is the intermediate layer inhibitory weight, and the Y-axis is the excitatory weight. On the heatmap colorbar, the 0 value is the minimum possible reward-per-trial, while 1 is the maximum possible reward-per-trial. The dot on the colorbar indicates the performance of the Linear Network. Weight-configurations in decision regime I are outlined in white on the heatmap. Regime II weight-configurations are to the upper left of those areas, and regime III are to the right. The weight-configuration achieving the most reward is indicated by the black “X.” The left heatmap of (A) shows performance in the absence of environmental uncertainty (*σ_η_I__* = 0 Hz), while the right heatmap shows performance in the presence of high uncertainty (*σ_η_I__* = 1.5 Hz). (B) The optimal E/I weights shift as uncertainty increases. The X-axis is environmental uncertainty (*σ_η_I__*), the left Y-axis shows the optimal excitatory weight and the right Y-axis shows the optimal inhibitory weight. (C) The shift from regime I (tan) to regime III (sky blue, colors as in Figure 3C). The level of uncertainty is on the X-axis and the curvature of the fit to the indifference curve is on the Y-axis. (D) A measure of the change in reward-per-trial as a function of noise for a regime I (*J*_+_ = 0.34, *J*_−_ = −0.01), regime II (*J*_+_ = 0.36, *J*_−_ = 0.0) and regime III (*J*_+_ = 0.35, *J*_−_ = −0.10) weight-configuration. The magnitude of the slope indicates the susceptibility to uncertainty. (E) The slopes of lines fit in (D) for all weight configurations. The slope for the Linear Network is again indicated by the black dot.

We then measured how robust different weight-configurations of the Hierarchical Network were to changes in the level of uncertainty. For each weight-configuration, we fit a line to the proportion of maximum reward as the uncertainty level increased, as shown in Figure 4D. The magnitude of the slope was taken as the susceptibility, with a greater slope indicating that the weight-configuration was less robust to uncertainty changes. These slopes are shown in a heatmap on Figure 4E, with the slope of the Linear Network indicated again with a black dot on the colorbar. We found that the regime III areas, in addition to displaying greater performance with high uncertainty, were also more robust to changes in uncertainty levels.

### 2.4 Regime II Results from Nonlinear Summation, While Regime III Results from Max-like Operation

Having established that the configuration of E/I weights defines decision-making regimes, and that these decision-making regimes are optimal depending on the environment, we next examined the functional transformation taking place within an intermediate layer attribute-specific area, and we analytically investigated the conditions for which different regimes are optimal.

The phase planes of dynamical systems are a highly useful tool for analyzing the functional properties of decision networks (Wong and Wang 2006; Wong, Huk, et al. 2007). For an intermediate layer area, each combination of weight-configurations and input levels produced a unique phase plane. We calculated phase planes for all combinations of an intermediate layer area’s excitatory weights (0.30 nA to 0.40 nA, spaced 0.05 nA) and inhibitory weights (0.0 nA to −0.10 nA, spaced 0.05 nA), and all input combinations of 0 to 40 Hz, spaced 0.5 Hz. This produced 793,881 total phase planes. For each phase plane, we computed a noiseless trajectory starting near the origin. The endpoint of that trajectory was taken as the output of the specific combination of network structure and inputs. The transformation of an input to output for different weight-configurations is shown in Figure 5A, with the input on the left and the outputs of three different weight-configurations on the right. We used these fixed point trajectory endpoints to approximate *T*_*A*,1_, *T*_*A*,2_, *T*_*B*,1_, and *T*_*B*,2_. The decision was determined by linearly summing these *T* values to obtain *C_A_* and *C_B_*, then choosing the maximum of these two choice values.

**Figure 5:**
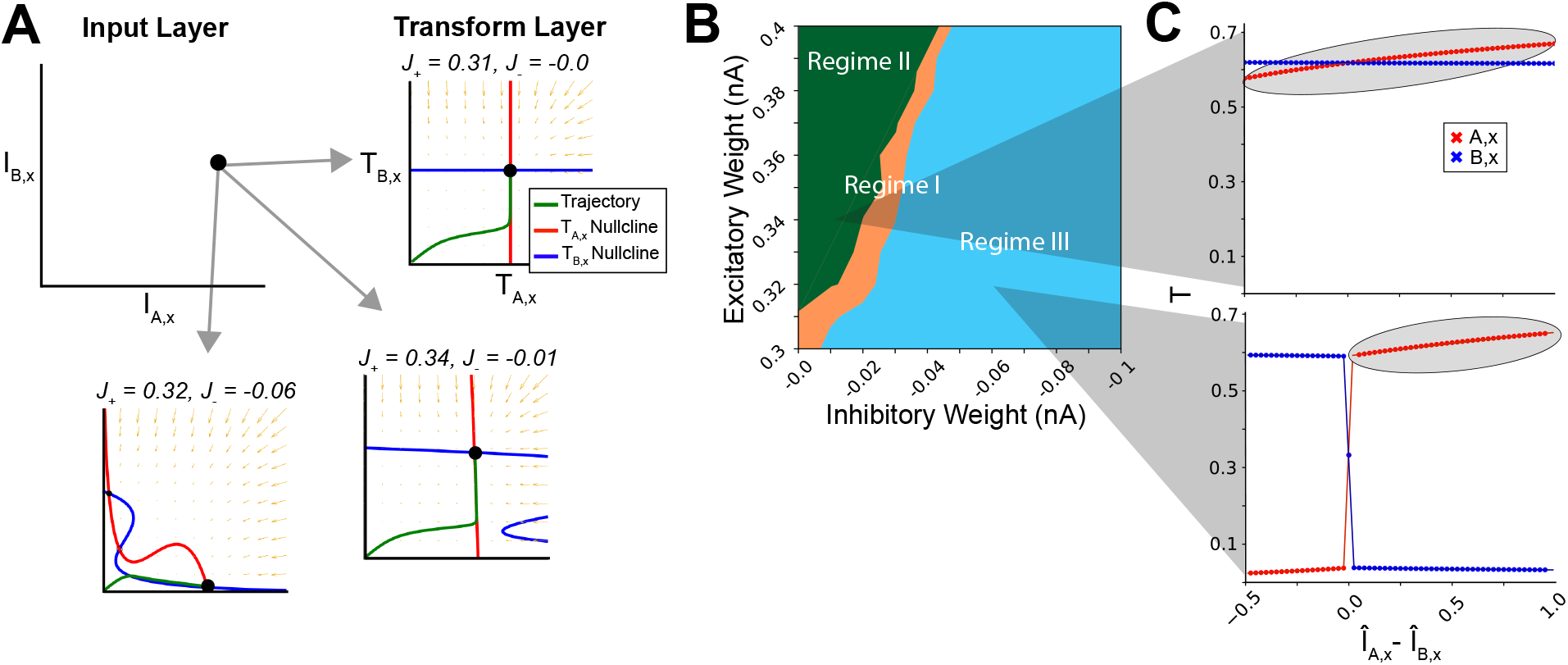
Functional Properties of Hierarchical Network Decision Regimes. *I*: input; *Î*: input normalized by the maximum input value; *T*: the transformed output of an intermediate layer; *J*_+_: excitatory weight; *J*_−_: inhibitory weight; red: choice alternative *A*; blue: choice alternative *B*; green: trajectory from a starting point of (0.06, 0.06); solid circles: stable fixed points; large solid circle: trajectory endpoint. (A) The input to an intermediate layer area is transformed to a fixed point output. The input values of *I_A,x_* = 15 Hz and *I_B,x_* = 10 Hz are presented graphically as coordinates on the left. These inputs were fed into intermediate layer areas (arrows) with unique weight-configurations. The resultant phase portraits of the transform layer are shown on the right and below for three weight-configurations. (B) The weight-space was partitioned according to the decision regimes arrived at through fixed point approximations. Indifference curves were computed for each weight-configuration by transforming offer input values using the endpoints of their phase-portrait trajectories, summing them to compute choice values *C_A_* and *C_B_*, then taking the larger of those values. The color scheme is the same as Figure 3C. (C) The input-output of a single attribute value as it is varied (*I_A,x_*) while the other (*I_B,x_*) is fixed for a regime II weight-configuration (top) and a regime III weight-configuration (bottom). The difference between the normalized fixed (*Î_B,x_*) and the varied value (*Î_A,x_*) is on the X-axis. The *T* value of the endpoints is on the Y-axis. Individual *T_A,x_* (red) and *T_B,x_* (blue) values are plotted, along with lines joining them to aid visualization.

Though the intermediate layer outputs to the choice area in the full dynamical model often did not reach the fixed points (due to the decision threshold), the fixed points did a reasonable job approximating behavior. To measure this, for each weight-configuration we fit a sigmoid to the choice behavior of the full model as a function of the difference between choice alternative fixed points (see Methods). Supplementary Figure S2 shows the distribution of *R*^2^ values. As over 90% of weight-configurations had an *R*^2^ > 0.70, we felt confident in use of fixed points as an approximation for analysis of intermediate layer function.

We then used fixed points to compute indifference curves, similar to those shown in Figure 3B. In Figure 5B, we show the decision-making regimes as computed from the intermediate layer fixed points (as in Figure 3C). Its qualitatively similar form to that of Figure 3C further supports that the fixed points provide a reasonable approximation of the full network.

Confident in the representativeness of the intermediate layer fixed points, we used them to investigate the input-output relationship of various intermediate layer weight-configurations. To do this, we first looked at how the output changes as one input is increased while all else is held constant. We then assessed the how multiple attributes interact as they pass through the intermediate layer.

To investigate how output changes as only one input is varied, we kept one of the attribute inputs (*I_A,x_*) to an intermediate layer area fixed, while increasing the level of the other attribute input (*I_B,x_*). The resultant *T_A,x_* and *T_B,x_* for a region II weight-configuration (*J*_+_ = 0.34 nA, *J*_−_ = −0.01 nA), and a region III weight-configuration (*J*_+_ = 0.32 nA, *J*_−_ = −0.06) are shown in Figure 5C. Note that in regime II (Figure 5C, top), the constant-input does not change as the magnitude of the variable input increases. Furthermore, the output level of the varied-attribute (outlined in gray) in regime II shows a decreasing rate of change as its magnitude grows. The same plot for regime III is shown in Figure 5C, on the bottom. In regime III, when the variable input is less than the fixed input, the output-value of the variable input is suppressed; however, when the variable input is greater than the fixed input, the output-value of the constant input is suppressed, while the output-value of the varied input grows approximately linearly. Thus, in region I and II, each input value passes through the intermediate layer whose degree of linearity is defined by the weight-configuration (Supplemental Figure S3A). In region III, however, a qualitatively different function is implemented such that the smaller input value is suppressed by a max-like operation (Supplemental Figure S3B).

Having established that different decision-making regimes maximize reward at different uncertainty levels, and then how the network produces the regimes, the natural question arises as to why these regimes are optimal in different environments. To answer this question, we turned to a mathematical analysis of simplified non-dynamical models. One model represented the extreme case of sequential max-operations (corresponding with regime III) and the other model that of a linear sum (regime I). We found that when the two choice alternatives are extremely dissimilar, sequential max operations can lead to sub-optimal choices. However, when the choice alternatives are similar, the max operations allow a network to improve the signal relative to noise (see Supplemental Information for proofs).

### 2.5 Choice Patterns of Those With Autism will Differ from Neurotypicals as Environment Uncertainty is Varied

Having shown that the Hierarchical Network can adopt multiple decision-making regimes, that the regimes are optimal in different environments, and how E/I tone defines regime-specific attribute transforms, we next hypothesized a framework where E/I tone can be modulated according to the degree of environmental uncertainty. We then applied this framework to investigate altered decision-making in ASD.

ASD is a developmental disorder with strong evidence for inhibitory dysfunction (Foss-Feig et al. 2017; Lee, Lee, and Kim 2017; Zikopoulos and Barbas 2013). Although E/I alterations observed in animal models of ASD are highly diverse, one of the more consistent findings is alterations involving parvalbumin positive (PV) interneurons, which have a strong influence on the overall inhibitory tone of a circuit. Gogolla et al. found that PV interneurons were reduced across several areas of cortex and the hippocampus in a diverse set of ASD mouse models (Gogolla et al. 2009). This selective reduction in PV interneurons was similarly found in the cortex of human postmortem tissue, where only the proportion of PV interneurons was decreased relative to controls (Hashemi et al. 2017). Decreases in PV populations are behaviorally relevant for ASD. For example, mice specifically engineered for PV depletion display phenotypes strongly liked with ASD, such as abnormal reciprocal social interactions, impairments in communication, and repetitive and stereotyped patterns of behavior (Wöhr et al. 2015). These and other findings have provided strong support for the “E/I imbalance,” theory of autism, where inhibitory tone is restricted relative to excitatory tone (Foss-Feig et al. 2017). Thus, manipulations of E/I tone in the hierarchical model can provide an avenue for linking the biology of ASD with behavior.

In addition to restricted inhibitory tone in ASD, there is evidence for a general intolerance of uncertainty (Boulter et al. 2014; Fujino et al. 2017; Vasa et al. 2018), and altered performance with environmental uncertainty (Milne et al. 2002; Spencer et al. 2000). Furthermore, differences have been found in multiattribute decision tasks (Foxe et al. 2015; Zaidel, Goin-Kochel, and Angelaki 2015). We therefore created a multi-attribute decision task with varying uncertainty as to the true value of attributes. The task, designed for pediatric subjects, involves choosing between baskets of food for “Harry the Hippo.” Harry’s favorite foods are M&Ms and Jelly Beans, which he enjoys equally. On a given trial, the network was presented with two baskets to feed Harry, each composed of M&Ms and Jelly Beans (Figure 6A). In the figure, quantities of the attributes are indicated by a horizontal lines on a vertical bars associated with each attribute. The goal over the course of the session is to feed Harry as much food as possible. Uncertainty is created by obscuring the exact position of the line with a yellow bar (right trial in Figure 6A); the wider the yellow bar, the more uncertain the attribute’s true value.

**Figure 6:**
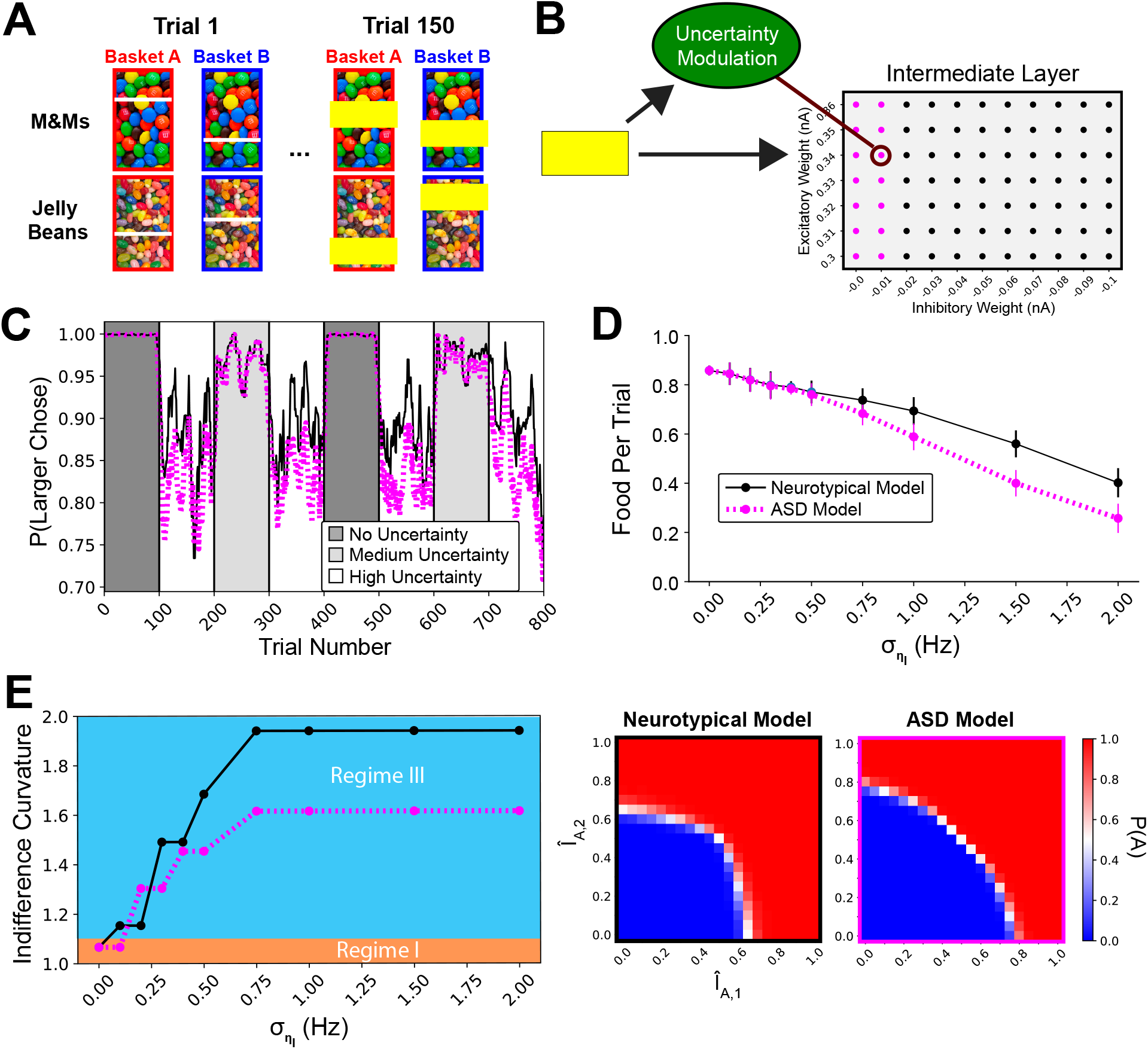
Task Simulation with Neurotypical and ASD Models. (A) The two-alternative, two-attribute task with varying levels of uncertainty. Alternatives, consisting of the attributes M&Ms and Jelly Beans, are simultaneously presented. The offer value of an attribute is indicated by the position of the horizontal white line on the vertical rectangle such that the higher the line, the greater the quantity of the attribute. Environmental uncertainty is controlled through a yellow bar obscuring the exact location of the white line (right trial). (B) A module was added (forest green) that modulated the intermediate layer E/I tone according to the uncertainty level. For the ASD model, inhibitory tone was restricted to a maximum magnitude of −0.01 nA, as is shown by the magenta dots in intermediate layer diagram. (C) A single session of the simulation, where the trial number is given on the X-axis and the proportion of trials where the larger option was chosen given on the Y-axis. The smoothed value for the neurotypical and ASD performance is given in black and magenta respectively. Blocks with no environmental uncertainty are in dark gray, medium uncertainty in light gray, and high uncertainty in white. (D) The reward-per-trial as a function of uncertainty for both the neurotypical and ASD models, with error bars drawn from sets of sessions. (E) The optimal indifference curvature as a function of uncertainty for the neurotypical and ASD models. To the right are example indifference curves for neurotypical (left) and ASD (right) at their maximum level of concavity. Input values (*Î*) are normalized by the reference choice alternative values’ linear sum.

The E/I tone of a brain area is not necessarily fixed. Indeed, there are several mechanisms by which the tone can be temporarily modulated (Zucker 1989; Semyanov et al. 2004; Avery and Krichmar 2017). The cholinergic diffuse modulatory system, in particular, provides a plausible mechanism for the source of E/I tone shift due to expected environmental uncertainty (Yu and Dayan 2005).

To allow the network to adapt to uncertainty levels, we added a module that used a look-up table to identify the optimal intermediate layer weight-configuration for a given environmental uncertainty level, then shifted weights in the attribute-specific intermediate layers to that configuration (green in Figure 6B). The neurotypical model was able to explore the full range of possible weights. The ASD model was implemented by limiting the possible inhibitory tone in intermediate attribute-specific areas to a maximum magnitude of −0.01 nA.

Sessions were composed of trial blocks with variable levels of uncertainty. A single session is presented in Figure 6C, with the smoothed proportion of the larger value chosen across trials shown for the neurotypical (black) and ASD (magenta) models. Note that during high-uncertainty blocks (white), one can already see a separation of performance levels. Analyzing food-per-trial as a function of uncertainty, we found that the ASD model did not differ from the neurotypical model when the uncertainty level was low, but began to diverge as uncertainty increased (Figure 6D). We also examined the change in the indifference curve as a function of uncertainty. The trend is shown in the left portion of Figure 6E, while specific instances of neurotypical and ASD indifference curves at the highest level are shown to the right. We found that there was a ceiling to the concavity of the ASD model’s decision behavior. Thus, the model makes two strong experimental predictions: 1) performance will be identical at low uncertainty, while subjects with ASD will show a greater fall-off in total reward as uncertainty increases; and 2) at high uncertainty, subjects with ASD will continue to show more linear-weighting of attribute values than neurotypicals, who shift to attending to the larger of the attribute values.

## 3 Discussion

We showed that a hierarchical network of areas defined by E/I tone is capable of performing a multi-attribute decision task, and can adopt behavioral regimes that maximize reward in a variety of environments. In environments where attribution values are certain, the network maximizes reward by adopting a regime that sums a linear translation of offer attribute values (regime I). However, as the environment becomes more uncertain, the network maximizes reward by implementing a max-like function to filter noise (regime III). We then proposed a framework where E/I tone can be modulated according to uncertainty. We used this framework to predict that, due to an imbalance of inhibition relative to excitation, subjects with ASD will diverge in behavior from neurotypicals as environmental uncertainty increases.

There is support in the literature for both the hierarchical and parallel structure of our model. Several prior computational models have proposed a similar parallel processing of attributes that are then joined in areas more proximal to the final decision (Roe, Busemeyer, and Townsend 2001; Balasubramani, Moreno-Bote, and Hayden 2018; Hunt, Dolan, and Behrens 2014; Cisek 2006; Christopoulos, Bonaiuto, and Andersen 2015). Furthermore, recent studies involving humans directly examined attribute-specific processing prior to integration in downstream areas. (Berker et al. 2019) looked at the representation of quantity and quality attributes in gift cards. They found that distinct areas were independently specific for each attribute, and that a third area showed activity correlated with the combined signals from the earlier areas. The importance of later-areas for the integration of value signals was demonstrated by (Pelletier and Fellows 2019) in patients with damage to the OFC and vmPFC. They found that these patients were able to make decisions based on single attributes (the form of elements in an object versus the configuration of elements) no different from controls. However, when patients had to make decisions based on integration of the attribute signals, they showed deficits. These results are expected from a neural system with parallel structure that converges for integration of attribute information.

A down-stream area providing a common representation of attribute value information is a key component of our model. Several electrophysiology studies have specifically investigated the kind of two-alternative multi-attribute choices that we have examined, while recording from down-stream areas relevant for decisions such as the OFC or ACC. Reward magnitude, probability, information value, social hierarchy and juice type have all been found to be represented in these areas, and in a manner where all attribute signals were available for decision-making (Hunt, Malalasekera, et al. 2018; Pastor-Bernier, Plott, and Schultz 2017; Pastor-Bernier and Cisek 2011; Blanchard, Hayden, and Bromberg-Martin 2015; Cai and Padoa-Schioppa 2014; Raghuraman and Padoa-Schioppa 2014; Cai and Padoa-Schioppa 2012; Xie and Padoa-Schioppa 2016; Padoa-Schioppa and Assad 2006; Padoa-Schioppa 2009; Munuera, Rigotti, and Salzman 2018; Pastor-Bernier, Stasiak, and Schultz 2019). However, it is important to recognize that the stimuli used to signify attribute information in these studies were unambiguous, and that NHP subjects were over-trained on the use of attributes to optimize reward. Thus, these studies all take place in the “low uncertainty,” range of our results. A quasi-linear attribute value transmission is therefore expected. Our results suggest that if environmental uncertainty is introduced, one should observe an over-representation of the larger attribute value in areas such as the OFC or ACC, and that the activity will correlate with animal behavior. This is a distinct prediction, highly feasible for experimentation.

Though the biology of how E/I tone of specific areas can be tuned according to task demands remains to be fully characterized by experiments, diffuse modulatory neurotransmitter systems provide the most plausible mechanism. Dopamine, norepinephrine, serotonin, nitric oxide and acetylcholine (ACh) have all been shown to influence the E/I tone of the cells that they target (Avery and Krichmar 2017). These influences can be complex, with cell-type specific shifts in excitation or inhibition implemented via a variety of mechanisms (metabotropic receptors, ion channels, etc.). The ACh system in particular displays the area-specific targeting (Zaborszky et al. 2015), the modulation of inhibitory tone (Disney, Aoki, and Hawken 2012; Picciotto, Higley, and Mineur 2012; Herrero et al. 2008; Kang, Huppé-Gourgues, and Vaucher 2014; Phillips et al. 2000; Sarter, Lustig, Howe, et al. 2014; Sarter, Lustig, Berry, et al. 2016; Thomsen, Sørensen, and Dencker 2018; Disney and Aoki 2008; Disney, Domakonda, and Aoki 2006; Disney, Alasady, and Reynolds 2014; Disney and Reynolds 2014) and the activity associated with known uncertainty (Monosov, Leopold, and Hikosaka 2015; Voytko et al. 1994; Gill, Sarter, and Givens 2000; Dalley et al. 2004; Marshall et al. 2016) that is required by our model. However, it remains to be investigated precisely how the brain can identify an task-optimal E/I tone for a circuit and then shift the circuit to that value.

Our work makes additional strong predictions for decisions of ASD subjects performing a multi-attribute decision task as uncertainty is varied. There are a number of existing experimental results in the field of ASD research that suggest our predictions are reasonable and worth investigating. Several studies show that when attribute information is clear, performance between ASD and neurotypical subjects is identical, while it differs in the face of noise or uncertainty. Dot field motion tasks, in particular, have been used to study these questions in subjects with ASD. The general finding has been that when coherence of the moving dot field is high (uncertainty is low), subjects with ASD and neurotypicals show similar performance (Milne et al. 2002; Spencer et al. 2000; Zaidel, Goin-Kochel, and Angelaki 2015). Indeed, in detection of velocity, and some coherence studies, subjects with ASD outperform neurotypicals (Chen, Norton, et al. 2012). However, when coherence levels decrease (uncertainty increases), subjects with ASD consistently show significantly greater decreases in performance. Similar effects of environmental uncertainty have been shown in ASD multi-attribute studies (Zaidel, Goin-Kochel, and Angelaki 2015; Foxe et al. 2015). Our results suggest a plausible mechanism for the effects observed in prior studies, while also proposing a specific experiment with falsifiable findings.

In this paper, we used a hierarchical neural network to investigate how biophysical properties, such as parallel structure and E/I tone, shape multi-attribute, multi-alternative decisions. This investigation produced strong predictions for electrophysiology experiments, as well as experiments involving human behavior. Our work thus suggests a new avenue for research connecting neural circuits to multi-attribute economic choice and to deficits associated with autism.

## 4 Acknowledgments

This research was partly supported by the National Institutes of Health (NIH) grant 062349, and the Simons Collaboration on the Global Brain program grant 543057SPI To XJW.

## 5 Methods

### 5.1 Model Architecture

#### 5.1.1 Cortical Areas

Areas in the networks were composed of mean-field attractors that have been used to model working memory, perceptual decision-making, as well as to model individual areas in multi-area models of cortex (Wong and Wang 2006; Wong, Huk, et al. 2007; Murray, Jaramillo, and Wang 2017; Mejias et al. 2016). Each area consisted of two populations, such that *c* = *A*, *B*. The dynamics of a given population is described by a single synaptic gating variable representing the fraction of activated N-methyl-D-aspartate receptor (NMDA) conductance and governed by the equation (Wong and Wang 2006),

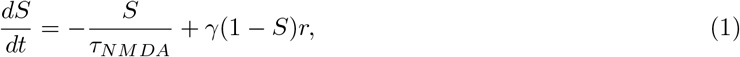

where the NMDA time constant *τ_N M DA_* = 60 ms, and the rate of saturation is controlled by *γ* = 0.641. The firing rate *r* was a function of the input current *I*, as defined by the curve relation first described in (Abbott and Chance 2005):

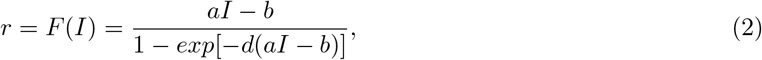

where *a* = 270 Hz/nA, *b* = 108 Hz, and *d* = 0.154 seconds. The total synaptic current consisted of recurrent (*I_rec_*), noisy (*I_noise_*), background (*I_o_* = 0.3297 nA) and external components such that,

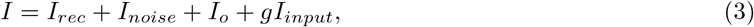

where the coupling constant *g* = 0.0011 nA/Hz converts the firing rate *I_input_* to current. As described in section 5.1.2, specific values of *I_input_* are notated as *I*_*A*,1_, *I*_*A*,2_, *I*_*B*,1_, or *I*_*B*,2_. For a given population *c* in area *n* of the the network, the recurrent current was given by the equation,

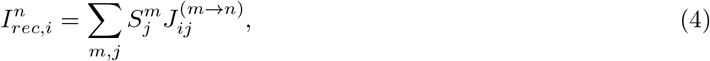

where 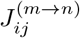 is the strength of the connection from population *j* in area *m* to population *i* in area *n*. The use of this equation in the Linear Network and Hierarchical Network is described in the following sections.

The noise currents for each population were defined by an Ornstein-Uhlenbeck process:

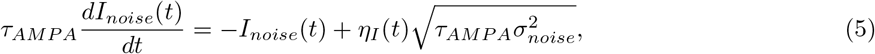

where the time constant *τ_AMPA_* = 2 ms, and *η_I_* is a Gaussian white noise term with mean zero, and variance 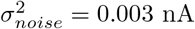.

#### 5.1.2 Inputs

External inputs were provided as firing rates, which could span the range of 0 to 40 Hz. Notationally, they are given by *I_input_*, such that the input of attribute 1 for choice alternative *A* was written *I*_*A*,1_, etc. An unspecified choice alternative is indicated with the population subscript “*c*” and and unspecified attribute by the subscript “*x*.”

When varying environmental uncertainty, a value *η_I_* was randomly drawn from 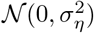 independently for each attribute on each trial, such that *Ĩ_c,x_* = *I_c,x_* + *η_I_*. The standard deviation *σ_η_I__* was used to control the amount of environmental uncertainty in a given block of trials.

In several figures, we refer to a normalized value of the input, *Ĩ_c,x_*. In those cases, the quantity is specified with respect to which the input is normalized.

#### 5.1.3 Linear Network

The Linear Network consisted of two layers. The first layer provided inputs from the individual attributes, in the form of firing rates *I_input_*. The attribute inputs to a given population were linearly summed, such that

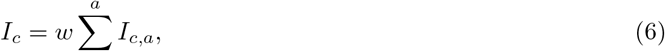

where *w* = 0.5. These values were then fed into the choice area described in section 5.1.1. As there was only one area, equation 4 simplified such that,

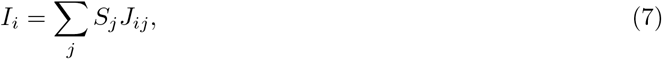

where for *i* = *j*, the connection strength was 0.3725 nA, and where *i* ≠ *j* the connection strength was −0.1137 nA (Wong and Wang 2006). The decision was determined when one of the choice area populations passed the firing rate threshold of 35 Hz.

#### 5.1.4 Hierarchical Networks

The input layer to the Hierarchical Networks was also in Hz, and specified by *I_c,a_*. In the Hierarchical Network, however, inputs were segregated by attribute and fed into an intermediate transform layer, consisting of a two-population area specific for each attribute. These areas were structured along the lines described in section 5.1.1.

In the intermediate layer areas of the Hierarchical Network, equation 4 was defined as follows. *J*_+_ controlled the strength of the excitatory connection of a population to itself (*A* → *A*, etc.), and *J*_−_, the strength of the inhibitory connection from one population to another (*A* → *B*, etc.). The strength of *J* between areas in the intermediate layer areas was set to 0. Connectivity from the intermediate layer (IL) areas to the choice layer (CL) was restricted to excitation between populations with the same selectivity, such that 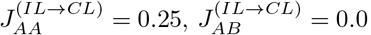, etc.

The excitatory and inhibitory weights in this intermediate layer (*J*_+_ and *J*_−_) were parametrically varied to define the specific Hierarchical Network weight-configurations described in the Results section. For simplicity, weights in all areas in the processing layer were symmetrically varied, such that when the recurrent excitation or cross-inhibition were changed, all areas in the processing layer assumed the new values. The outputs from this processing layer were dynamically fed into a choice area of the same type as the choice area of the Linear Network in section 5.1.1. To clarify notation, when an input *r* is passed through an intermediate transform layer area, its associated transformed output is referred to as *T*. The value *T* is that of the synaptic gating variable from equation 1.

### 5.2 Model Behavior

#### 5.2.1 Psychometric Curves

For the psychometric curves shown in Figure 2, a single choice alternative attribute was varied while all others were held constant. 1,000 trials were run for the Linear Network, and for the Hierarchical Network. We then calculated the percent of those trials where the networks chose the option associated with the varied attribute. To these points, we fit a sigmoid of the form:

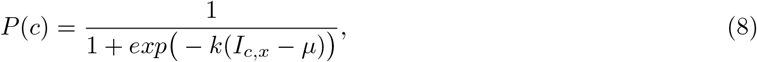

where *P*(*c*) represents the proportion of the time the varied choice alternative was chosen, *k* dictates the slope of the sigmoid, *I_c,x_* indicates the value of the varied attribute, and *μ* provides the centering of the sigmoid.

#### 5.2.2 Indifference Curves

The indifference curves of Figure 3C and 5B were calculated by holding the values for choice alternative *A*(*I*_*A*,1_ and *I*_*A*,2_) constant as a reference, while independently varying *I*_*B*,1_ and *I*_*B*,2_. The reference attribute inputs were both 20 Hz, and the varied attribute inputs ranged from 0 Hz to 40 Hz. All possible combinations of the varied attributes were then simulate for 1,000 trials, where the networks chose between the varied alternative and the reference. This was done for the Linear Network, as well as for several processing-layer weight-configurations of the Hierarchical Network. The processing-layer weight-configurations were all possible combinations of the recurrent excitatory weights 0.30 to 0.40 nA, spaced 0.01 nA with the cross-inhibitory weights 0.0 to −0.1 nA, spaced 0.01 nA. Indifference values were then taken to be those where the network chose evenly between the alternatives.

To fit an indifference curve, we first normalized the indifference values such that the minimum in either direction was 0, and the maximum was 1. We then fit the equation,

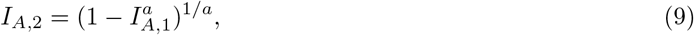

where *a* was used to define the shape of the indifference curve. If *a* < 1, the curve was classified as convex. If *a* = 1, it was classified linear, and if *a* >1 it was classified as concave. These classifications were then used in Figures 3C and 5B to partition the weight space into areas that produced convex, linear or concave indifference curves.

#### 5.2.3 Reward-Per-Trial

To simulate a choice experiment, we computed the choice behavior of the network in response to *I_c,x_* values in all possible combinations of the inputs ranging from 10 to 20 Hz, with a step size of 1 Hz, producing 630 possible combinations. This was done for the Linear Network, as well as for the Hierarchical Network with all combinations of the recurrent excitatory weights ranging from 0.30 nA to 0.36 nA, spaced 0.01 nA, with the cross inhibitory weights ranging from 0.0 nA to −0.1 nA, spaced 0.01 nA. Each choice was run for 1,000 trials, from which the percentage chosen was calculated.

Reward was considered to be the total value of the attributes for the chosen alternative, regardless if it was the larger of the two. Thus, on each trial there was a maximum reward (the larger choice alternative) and a minimum reward (the smaller choice alternative). The reward-per-trial was computed by simply averaging the reward received across all trials. We did the same for the minimum and maximum reward. We then set 0 to be the minimum reward-per-trial and 1 to be the maximum reward-per-trial. Those values were used to bound the colorbar in Figure 4A.

#### 5.2.4 Uncertainty Robustness

To compute the robustness to uncertainty, as shown in Figure 4E, we systematically stepped the *σ_η_I__* as described in section 5.1.2 through the values 0 to 0.5 Hz with a step size of 0.1, as well as 0.75 to 1.0 Hz with a step size of 0.25, and 1.5 to 2 Hz with a step size of 0.5. For the Linear Network and each weight-configuration, the the reward-per-trial was calculated at each of these uncertainty levels. A linear function was then fit to the reward-per-trial as a function of uncertainty level of the form,

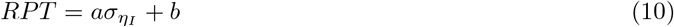

Where *RPT* is the reward-per-trial, *a* is the slope and *b* the intercept. The robustness to uncertainty was assessed using the magnitude of *a*, such that the greater the magnitude, the more susceptible.

### 5.3 Fixed Point Analysis

To compute the fixed points and phase portraits of a processing layer area, we used the python package pydstool (Clewley 2012), parameterized by the equations described in 5.1.1, with the internal noise was set to 0. We first defined an attribute input as a coordinate in euclidean space space whose positions are defined by the level of each input (Figure 5A, left). We then defined a phase plane by applying these inputs to each weight-configuration. A starting position in each phase plane was selected near the origin at (0.06, 0.06), and a noiseless trajectory was computed. The end point of that trajectory was taken as the output of the processing layer.

#### 5.3.1 Calculation of Convexity

The trajectory endpoints were computed for all inputs and weight-configurations described in section 5.2.2. These endpoints were treated as the values *T*. They were then linearly summed to determine the value of a choice alternative, such that,

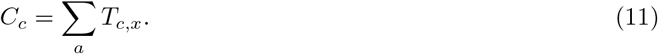

The indifference point was taken to be when the *C_A_* ≈ *C_B_*. Convexity was then computed and the weight space was partitioned using the methods described in section 5.2.2.

#### 5.3.2 Calculation of Functional I/O: Linearity

To determine the functional transformation of inputs as they passed through the processing layer, for each weight-configuration we kept *I_B,x_* constant at 20 Hz while increasing *I_A,x_* from 0 to 40 Hz. We then recorded the resultant *T_A,x_* and *T_B,x_* as the output of the functional transformation. We identified when the transition of *T_A,x_* stabilized and fit the function,

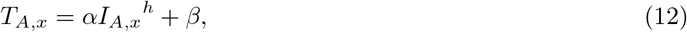

where *h* was used to determine the linearity of the transformation. An *h* ≈ 1 indicated that the network produced an output that was a linear function of the input. If *h* < 1 however, the network produced a sub-linear output. The value of *h* was plotted in the heatmap of Figure S3A.

#### 5.3.3 Calculation of Functional I/O: Interaction

To calculate the full interaction of values, we computed *T_A,x_* as we varied both *I_A,x_* and *I_B,x_* from 0 to 40 Hz, with a step size of 0.5 Hz. We then plotted the values of *T_A,x_* shown in the heatmap of Figure S3B.

### 5.4 Neurotypical and ASD Models

We introduced a module that used a look-up table to assess the level of uncertainty in the environment, and then set the weight-configuration in the intermediate layer areas to a tone that optimized performance. The neurotypical model was able to utilize the full range of excitation (0.30 nA to 0.40 nA) and inhibition (0.0 nA to −0.1 nA). The ASD model was able to utilize the full range of excitation, but the inhibitory tone was limited to a maximum of −0.01 nA. We then presented the models with blocks of trials at different uncertainty levels, and calculated the average reward-per-trial (section 5.2.3) during each block.

## Supplemental Information

### 1 Analysis of Linear Model and Sequential Max Model

The Hierarchical Network achieves maximum reward in an environment without uncertainty by adopting a weight-configuration that performed linear integration of attribute values (similar to the Linear Network), while in an environment with high uncertainty a max-like operation achieved maximum reward. In this section, we use simplified versions of the model to analytically investigate these observed phenomenon.

#### 1.1 The Linear Model And the Sequential Max Model

We will consider and compare two models of multi-attribute decisions. In the linear model (LM), attribute input values (*I*) for each choice alternative were linearly combined to reach a choice value *C_A_* or *C_B_*, and then the maximum of those values was taken as the choice (as in Figure S1A). The sequential max model (SM) consisted of a series of max operations (as in Figure S1B-C). These two models are extreme cases of decision regimes adopted by the Hierarchical Network, as described in the main body of the paper. Though we use the same notation as the main body of the paper for clarity (*I*, *C*, etc.), the values in this section are unitless.

**Figure S1:**
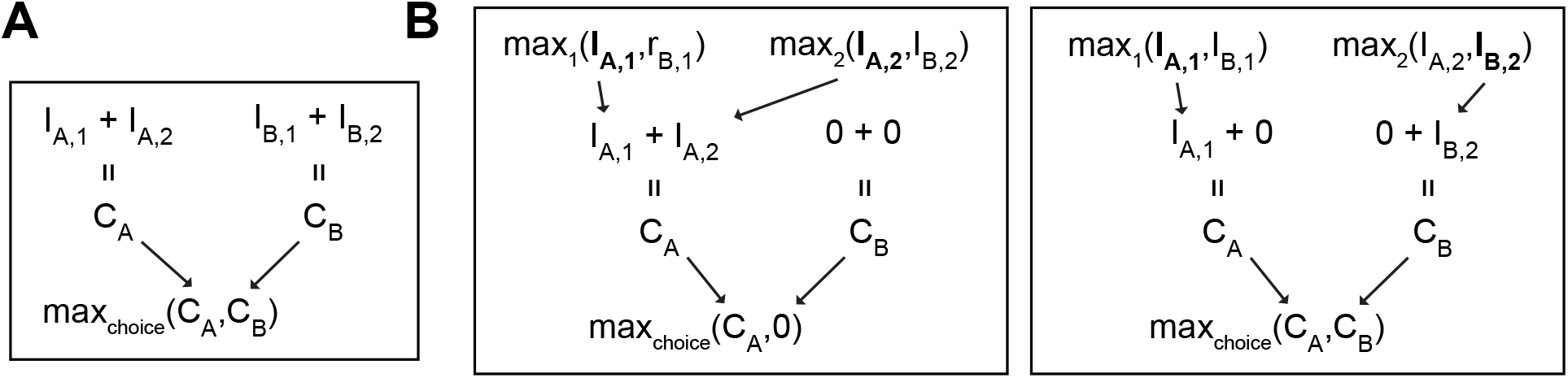
Static Decision Models. (A) The linear model, where attribute input values (*I*) are summed into a choice value (*C_A_* or *C_B_*). Choice is then determined with a max operation. (B) The sequential max model with a max operation at the attribute level and the choice level. The surviving values from the first max operation are summed to calculate *C*. Two examples of the sequential max model are shown, where the larger attribute input values are bolded. On the left, both attribute inputs associated with *A* are larger, and so these are used to calculate the *C_A_*. Since the *B* attribute inputs are smaller, they do not survive the first stage, and so *C_B_* = 0. On the right, an attribute associated with each choice alternative survives the first stage, and so both *C* values are nonzero.

Our analysis will focus on the most basic case, where there are two choices composed of two attributes. For notation, we will index the first choice as *A* and the second choice as *B*. Note that the number of choices could be expanded arbitrarily. We index the two attributes associated with each choice as 1 and 2. When the two attributes are interchangeable, it will be indicated with the subscript *x*. An attribute could be the color of a choice alternative, the monetary value, or (if it is edible), its flavor. As an example, the input to the model of the monetary value (1) associated with choice alternative 1 (*A*) could be written as *I*_*A*,1_. The flavor (2) associated with choice alternative 1 (*A*) could be written as *I*_*A*,2_. We can then write the initial offers as [*I*_*A*,1_, *I*_*A*,2_] and [*I*_*B*,1_, *I*_*B*,2_].

##### 1.1.1 The Linear Model: Equations

The LM was meant to capture the Linear Network, along with the case of the Hierarchical Network model performing a linear addition of the attribute values. The operations of the LM are shown in Figure S1A. In this model, the attributes associated with each choice alternative are first summed, such that,

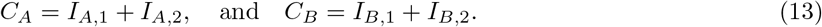

The second decision stage then takes the maximum of these choice values as the choice of the network, consisting of the operation,

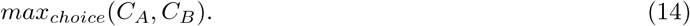

For the linear model, we will often thus simplify the decision stage by writing,

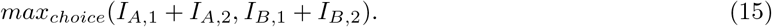

##### 1.1.2 The Sequential Max Model: Equations

The SM is named for the series of max operations performed, first at the input level, and then with the final decision. Those operations are shown in Figure S1B-C. This model produces the concave decision behavior described in the main text. The first stage of SM implements operations on the attribute level,

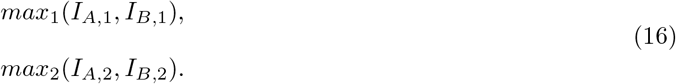

The key feature is that *only the maximum of that operation is passed onto the next level*. To gain intuition, we can walk through a couple of examples. First, consider the case where,

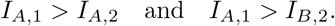

The decision process is shown in on the left of Figure S1B. Since only the winners of the first max operation (bolded in Figure S1) are passed to the next stage, the choice values will be,

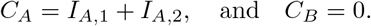

We can take another example to further illustrate the point (Figure S1C right). In this case, we are going to define the values such that,

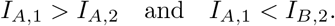

After these values pass through equation 16, the choice values are,

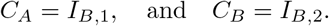

Because of this, the *C* values and composition will be specified in each examined case.

##### 1.1.3 When Offers are Dissimilar, Max Produces Irrational Decisions

Here we will show that, in an edge case, the LM will choose the larger of the option, while the SM paradoxically chooses the lower.

We will consider the edge case where the value of *A* is less than the value of *B*,

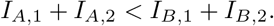

We will further constrain the values so that,

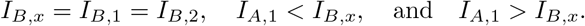

In the LM, by substituting the values into equation 15, we can specify the decision stage as the operation,

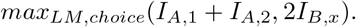

Given the constraints, the LM will choose *B*, which is the larger of the two offers.

In the SM, by use of equation 16, the first stage will consist of,

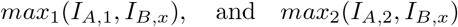

Again, given the constraints, after the first stage we will have,

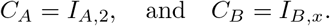

By use of equation 14, the decision stage will be,

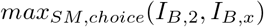

As *I*_*A*,2_ > *I_B,x_*, the SM will choose *A*, even though the total value of *B* is greater.

**Conclusion 1** When the offers are highly dissimilar, there will be cases where the SM will choose the offer with the smaller total value.

##### 1.1.4 Uncertainty in the Linear Model and Sequential Max Model

In a biological system, uncertainty from noise can arise externally via the stimulus or value representation, or internally via transmission between areas, or internally during the decision-making operation. For comprehensiveness, this section will consider how, at all stages, noise effects the LM and SM frameworks. Here, we are agnostic as to the source of the noise (stimulus, representational, transmission, background, etc.), and simply assume a random process, such that the noise term *η* is drawn from a uniform distribution with the limits [−*η_max_, η_max_*]. By using a uniform distribution, we can more easily delineate the extreme cases. When there are two stages at which the noise term is added, we will simplify their expression notationally by writing,

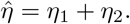

we will consider the two-attribute, two-alternative decision paradigm. To make the algebra clearer, we will introduce new variables *D*, *E* and scaling term *γ* such that,

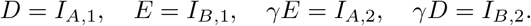

We will also stipulate that,

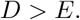

By maintaining the symmetry of the attributes across choice alternatives, we can finely control the magnitude differences, while keeping the choice alternatives similar to one another. This can be better understood by examining the scaling term *γ*, which tunes the magnitude of the difference between the choice alternatives.

For clarity, we will first write rewrite equation 14 using the values as,

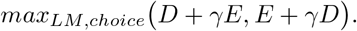

It is easy then to see that the larger C will be dictated by the value of *γ*. If *γ* > 1, then *B* will be larger, if *γ* < 1, *A* will be larger, and if *γ* = 1 the total value of each choice alternative will be equal. Adding a combined noise term, we arrive at,

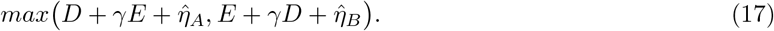

If we define *γ* as such that,

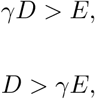

we can follow the algorithm, and arrive at the SM framework’s final decision stage where,

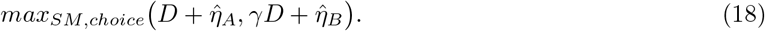

The challenge then is to understand the relationship between scaling term *γ* (which controls the absolute difference between choice alternatives) and noise term *η*.

##### 1.1.5 Signal to Noise in the Linear Model Scale with the Difference Between Attributes

We will show that the noise term will be irrelevant in decision making as long as,

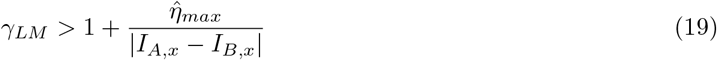

Rewriting equation 17 so that *A* is the winner we get,

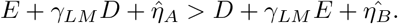

Giving alternative *B* the maximum benefit of noise we have,

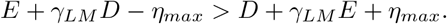

Solving for the *γ_LM_* terms on one side of the equation we get,

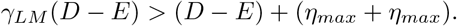

By defining

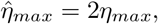

and dividing through by (*D* − *E*), and then back substituting the *I* values, we arrive at equation 19. It is important to see here that the minimum value of *γ_LM_* depends primarily on the term,

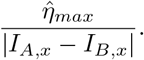

The smaller this term, the smaller *γ_LM_* can be.

**Conclusion 2** The minimum possible value of scaling term *γ_LM_* depends on the difference between *I_A,x_* and *I_B,x_*. Thus, the larger the difference between the attribute values, the smaller the difference need be between the choice alternatives.

##### 1.1.6 Signal to Noise in the Sequential Max Model Scales with Magnitude of the Larger Attribute

It can be shown that the value of *γ_SM_* needed to render the noise irrelevant is given by,

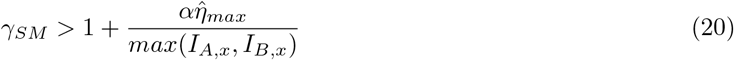

Now, it is reasonable to consider the possibility that in a SM, the additional operation creates an additional source of internal noise. We will specify its scale using a variable *α* where,

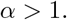

We will start by rewriting equation 18 so that *A* will be chosen over *B*. This can be specified by,

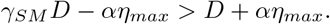

Solving for the *γ_SM_* terms the equation becomes,

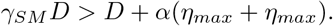

By dividing through by *D*, making the 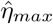 replacement, and back-substituting the *I* values, we arrive at equation 20. The important feature to note is that the minimum value of scaling term *γ_SM_* depends on,

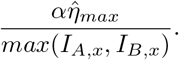

Unlike the LM, the denominator has only the larger of the attribute values.

**Conclusion 3** In the SM, the larger the largest attribute value relative to the noise, the less the magnitude difference between choice alternatives need be.

##### 1.1.7 When Offers are Similar and in the Presence of Noise, Sequential Max Improves Signal

We will show that,

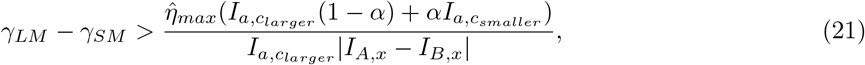

where,

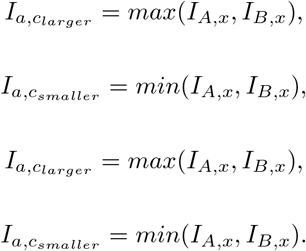

We start by subtracting equation 20 from equation 19.

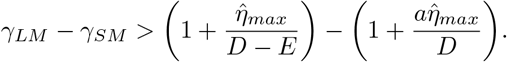

By expanding the right side of the inequality we get,

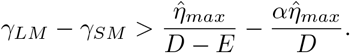

Putting both terms on the right side over the same denominator produces equation 21. Because *D* > *E* by definition, the denominator on the right will always be positive. However, for the inequality to hold, we need to consider what must be true of the values of *D*, *E*, *a*, *γ_LM_* and *γ_SM_*. By doing so, we will be able to draw some interesting conclusions regarding the two frameworks. First, lets consider the case where,

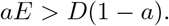

This means that the numerator on the right side of the inequality will be positive. Thus for equation 21 to be true,

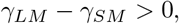

and therefore,

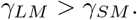

**Conclusion 4** When the two attributes are similar, it will require a greater difference between the choice alternatives (larger scaling term *γ*) for the *LM* model to recognize the more valuable choice alternative. If one compares equation 19 with equation 20, in equation 19, the noise is divided by |*I_A,x_* − *I_B,x_*|, while in equation 20 it is divided by *max*(*I_A,x_*, *I_B,x_*). In the LM, the closer the attribute values, the less the noise is reduced. In the SM, the smaller attribute value is no longer present; thus the noise term is always reduced. Furthermore, if we consider the total noise in both models to be equal by setting,

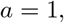

then the numerator simplifies to,

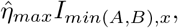

which is always positive. In such a scenario, conclusion 2 will always be true.

##### 1.1.8 If Internal Noise For the Sequential Max Model is Large, The Linear Model is Superior

Now, let us consider the extreme border-case where,

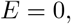

and thus,

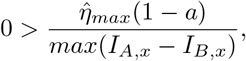

By itself this would say nothing about the value of *γ_LM_* – *γ_SM_* since it is possible that,

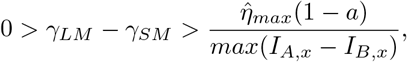

or,

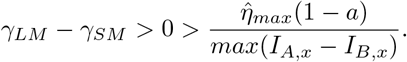

Yet, as goal is the find the minimum possible value of the *γ*s such that the inequalities hold, we can look back to equations 19 and 20 for insight. In the border-case, equation 19 becomes,

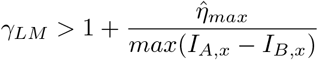

If minimize the value of *γ_LM_* that would keep the inequality true, and assume that *a* is sufficiently large, we get the inequality,

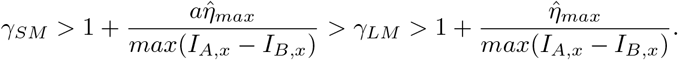

And thus,

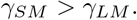

**Conclusion 5** Due to the additional noise caused by multiple SM neural operations, if the difference between the largest attribute and smallest attribute in each alternative is sufficiently large, the LM framework will require less of a difference between the alternatives in order to choose the greater choice alternative.

### 2 Supplemental Figures

**Figure S2:**
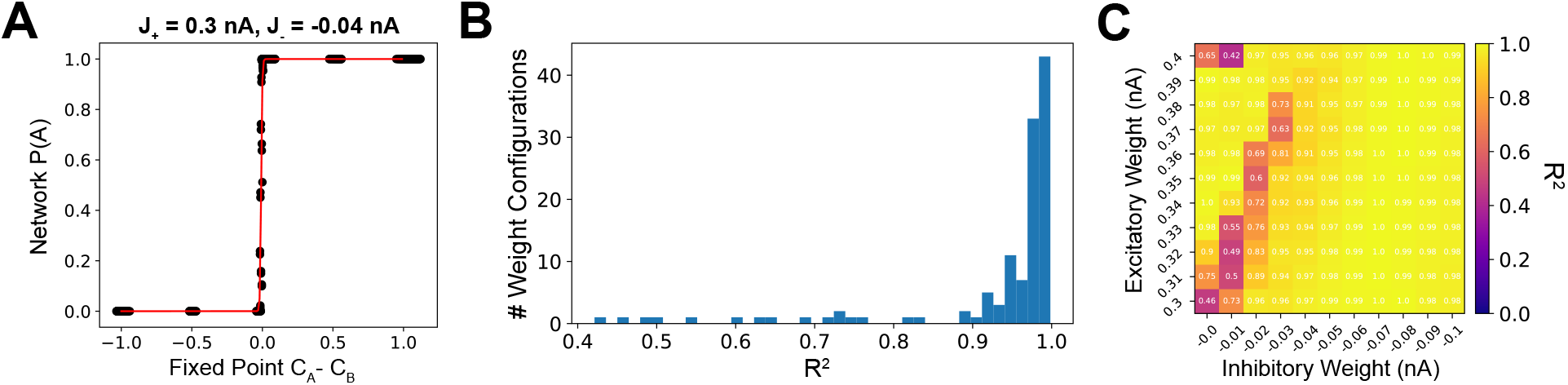
Intermediate Layer Area Fixed Points Fit to Full Network Behavior. Fixed point trajectory endpoints were used to approximate *C_A_* and *C_B_*. A sigmoid was then fit to the *P*(*A*) of the network as a function of fixed point *C_A_* – *C_B_* for each weight-configuration. The fit of a single weight-configuration is shown in A). An *R*^2^ value was then computed for each fit. A histogram of those *R*^2^ values is given in B), and in C) they are organized by weight-configuration.

**Figure S3:**
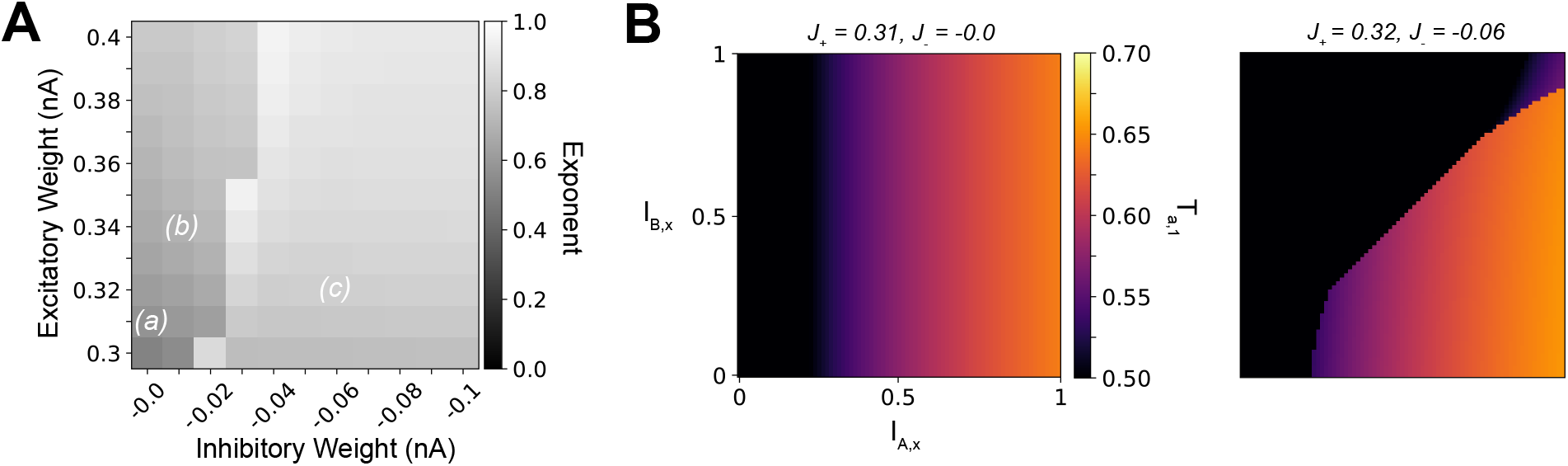
Functional Transformations by Hierarchical Network Intermediate Layer Areas. (A) Polynomial functions of the form *T*_*A*,1_ = *αI*_*A*,1_^*h*^ + *β* were fit to the constant-regions outlined in gray on Figure 5C. The value of the *h* exponent associated with each weight-configuration is indicated by the intensity of the heatmap. (E) The resultant *T_A,x_* as *I_A,x_* and *I_B,x_* are varied for a regime I (left) and a regime III (right) weight-configuration.

